# Establishing the foundations for a data-centric AI approach for virtual drug screening through a systematic assessment of the properties of chemical data

**DOI:** 10.1101/2024.03.28.587184

**Authors:** Allen Chong, Ser-Xian Phua, Yunzhi Xiao, Woon Yee Ng, Hoi Yeung Li, Wilson Wen Bin Goh

## Abstract

Researchers have adopted model-centric artificial intelligence (AI) approaches in cheminformatics by using newer, more sophisticated AI methods to take advantage of growing chemical libraries. It has been shown that complex deep learning methods outperform conventional machine learning (ML) methods in QSAR and ligand-based virtual screening^1–3^ but such approaches generally lack explanability. Hence, instead of developing more sophisticated AI methods (i.e., pursuing a model-centric approach), we wanted to explore the potential of a data-centric AI paradigm for virtual screening. A data-centric AI is an intelligent system that would automatically identify the right type of data to collect, clean and curate for later use by a predictive AI and this is required given the large volumes of chemical data that exist in chemical databases – PubChem alone has over 100 million unique compounds. However, a systematic assessment of the attributes and properties of suitable data is needed. We show here that it is not the result of deficiencies in current AI algorithms but rather, poor understanding and erroneous use of chemical data that ultimately leads to poor predictive performance. Using a new benchmark dataset of BRAF ligands that we developed, we show that our best performing predictive model can achieve an unprecedented accuracy of 99% with a conventional ML algorithm (SVM) using a merged molecular representation (Extended + ECFP6 fingerprints), far surpassing past performances of virtual screening platforms using sophisticated deep learning methods. Thus, we demonstrate that it is not necessary to resort to the use of sophisticated deep learning algorithms for virtual screening because conventional ML can perform exceptionally well if given the right data and representation. We also show that the common use of decoys for training leads to high false positive rates and its use for testing will result in an over-optimistic estimation of a model’s predictive performance. Another common practice in virtual screening is defining compounds that are above a certain pharmacological threshold as inactives. Here, we show that the use of these so-called inactive compounds lowers a model’s sensitivity/recall. Considering that some target proteins have a limited number of known ligands, we wanted to also observe how the size and composition of the training data impact predictive performance. We found that an imbalance training dataset where inactives outnumber actives led to a decrease in recall but an increase in precision, regardless of the model or molecular representation used; and overall, we observed a decrease in the model’s accuracy. We highlight in this study some of the considerations that one needs to take into account in future development of data-centric AI for CADD.

## Introduction

The discovery and development of a new drug and subsequently, bringing it onto the market is challenging and a major burden on time and finances. In a study of 63 drugs developed by 47 companies (between 2009 and 2018), it was estimated that the average cost of bringing a new drug to market is USD 1.33 billion^4^. To reduce cost and time, attention has shifted to using computer-aided drug design (CADD) methods, which has been relatively successful^5^. CADD allows researchers to better focus on drug experiments that have a higher likelihood of success.

In recent years, interest in AI/ML, particularly deep learning methods for CADD has grown. Deep neural networks have been shown to outperform machine learning (ML) algorithms like random forest (RF) and support vector machine (SVM) for quantitative structure-activity relationship (QSAR) and ligand-based virtual screening (LBVS). An example being Dahl et al.’s multi-task deep neural network (MT-DNN) which was the best performing architecture in the Merck Molecular Activity Kaggle Challenge^2,3^. A molecule is normally tested in multiple assays and their MT-DNN was trained so that the multiple output neurons each predicts the activity of the input molecule in a different assay. Following the Merck Challenge, researchers performed a detailed study that specifically compared the performance of deep neural nets (DNN) models to RF models and showed that DNN models routinely made better predictions on a series of large, diverse QSAR datasets generated as part of Merck’s drug discovery efforts^2^. However, contrary to findings of the above studies, when using DNN, RF and variable nearest neighbour (one of the simplest ML methods) to predict the molecular activities of 21 *in vivo* and *in vitro* datasets, Liu et al. reported that the overall performance of the three methods were similar^6^. This work leads us to question whether this trend to develop more advanced and complex AI methods would really result in any further significant improvements for QSAR/virtual screening.

In 2021, Dr Andrew Ng proposed that AI scientists should attempt to develop data-centric AI approaches instead of the currently pursued model-centric AI approaches^7,8^.

A model-centric AI approach involves improving/building on the AI algorithm to improve the model’s predictive performance ^8–10^. In contrast, a data-centric AI approach is one where automated methods are developed to improve data quality or to tune the data used by the predictive model – for example, such methods could check for consistency in the labelling of the data and automatically flag mislabelled data – and in so doing, improve the model’s predictive performance.

With some chemical databases like PubChem containing well over 100 million compounds^11^ and with the urgent need for large, clean datasets for deep learning methods, cheminformaticians are challenged to manually curate clean and consistent benchmark chemical datasets from large chemical libraries. Thus, it makes sense to develop data-centric AI approaches that can automatically do this for CADD. However, before we can develop such an approach, we need to understand what attributes/properties constitute good data for CADD. We believe there are four pillars of cheminformatics data that drives AI performance – namely, data representation, data quality, data quantity and data composition – and we were keen to investigate how each of these pillars contribute to an improved AI performance.

AI research generally begins with the extraction and acquisition of high quality data. But as data sets become larger and more sophisticated, manual data cleaning, restructuring and feature engineering becomes more challenging^12–14^. In a recent study, Northcutt et al.^15^ found label errors in the test sets of 10 of the most commonly-used computer vision, natural language, and audio benchmark datasets. On average, they found 3.3% errors across the 10 datasets - for example, the ImageNet validation set contained 6% (over 2,900 errors) label errors. Poor data quality can lead to biased AI models while insufficient data results in non-representative models that are unable to generalize and accurately predict from real-world data. However, in cheminformatics, it would appear that the latter is not an issue as there has been an exponential growth in chemical data: in 2010, PubChem contained 27,443,646 records of unique chemical structure compounds and by 2021, this had grown to around 111 million unique chemical structures (∼four-fold) ^16,17^.

Given Northcutt et al.’s findings, we felt that it was important for us to build a new benchmark chemical dataset that we could be confident of and which could be used with an AI to achieve superior performance. Only when we have a superior AI model that gives exceptional performance can we be certain that any change in the AI’s performance is indeed due to perturbations in our data and not because of an imperfect AI model. Thus, we carefully curated a new dataset of BRAF actives and inactives that we used for developing LBVS AI models for BRAF ligands. BRAF ligands are a well-studied class of drugs and there is much interest to develop potent BRAF antagonists that will suppress the actions of the constitutively-active mutant BRAF protein found in cancers like colorectal cancer and lung adenocarcinoma ^18–21^. Our results showed that that models trained on our BRAF dataset could attain near-perfect accuracy and so, this allowed us to design a set of experiments to test how each of the four pillars of cheminformatics data impacts on AI performance.

Enhancing data is not just about increasing the size (and representativeness) of the dataset but also about choosing the right data representation for a learning task. We tested the performance of different ML algorithms employing various molecular representations to evaluate how molecular representations affect ML algorithm performances. There have also been studies that have proposed the use of merged molecular representations^22–24^. This multi-representation of molecular information is powerful, and constitutes a form of multi-view learning^25^, an emerging direction in AI/ML with implications for improved generalization performance. We systematically tested if paired fingerprint combinations can do a better job of describing a molecule and outperform a single (standalone) fingerprint in LBVS. For this, we tested 10 standalone fingerprints and their 45 paired combinations to shed light on (1) what is the best type of molecular representation for virtual screening (i.e., substructure key-based, topological, path-based or circular fingerprints, and single or merged fingerprints) and (2) how the interplay between molecular representations and different ML algorithms contributes to the changes (if any) in predictive performance. In all, we developed and assessed 1,375 predictive models for LBVS of BRAF ligands.

Next, we investigated the impact that data quality, data quantity and data composition have on predictive performance. Using four top predictive models that utilizes only a single molecular fingerprint, namely, [i] SVM+ECFP6, [ii] RF+ECFP6, [iii] SVM+Daylight-like, and [iv] RF+Extended, we show how current conventional usage of data in CADD can be improved to enhance data quality. For example, a common practice in CADD is the use of DUD-E decoys as inactives but a recent study showed that a hidden bias in these decoys affects predictions for structure-based virtual screening (SBVS) ^28,29^. Thus, we wanted to see if this hidden bias also affects LBVS. In addition, we also explored how dataset size and composition impacts the performance of the AI models.

## Results

### Assessment of machine learning algorithms for ligand-based screening

Five balanced training datasets of BRAF actives and inactives were used to train five conventional machine learners: k-nearest neighbours (kNN), Naïve Bayes (NBayes), gradient-boosted decision tree (GBDT), RF and SVM. We created in total 1,375 predictive models - 5 predictive models for each ML algorithm using one of 55 different molecular representations. For all predictive models, we observe that no real difference between the average accuracy achieved in cross-validation and that achieved on the hold-out test set, for all 55 molecular representations used (Figure 1 and Supplemental Table 1); for example, for the SVM model trained with Estate fingerprint (the worst performing SVM model), cross-validation attained an accuracy of 85.2% while testing gave an accuracy of 85.7% and in some cases, the average accuracy achieved in cross-validation and testing were the same (SVM using either CATS2D [accuracy=97.1%] or Extended [accuracy=98.1%]). This suggest no overfitting, regardless of the ML algorithm or molecular representation used, and that our training and testing sets are representative of the BRAF class of drugs.

The best performance was achieved by an SVM model using the paired (ECFP6+Extended) fingerprints with an average percentage accuracy of 99.05% on the hold-out test set, followed very closely by the RF model using only the ECFP6 fingerprint that gave a 98.48% accuracy. SVM and RF had previously been shown to perform well in virtual screenings for BACE1 inhibitors when compared against other conventional machine learners (RF achieving AUC-ROC of 0.867)^26–28^ but to the best of our knowledge, the level of accuracy achieved here for our BRAF ligand dataset is unprecedented for a LBVS platform.

In general, all five ML algorithms tested were able to achieve similar levels of predictive performance, with the best performing models for each ML algorithm all achieving an accuracy of above 97%. We should also highlight that there was no feature selection performed on our training data prior to its use. In fact, the RF models were able to give a superior performance without even needing to tune the default parameters.

Among all predictive models, the worst performing model was the Naïve Bayes model using the Estate fingerprint (accuracy = 59.4%). But this shouldn’t come as a surprise. The conditional independence assumption of Naïve Bayes classifiers means that it can use high-dimensional features with limited training data compared to more sophisticated methods. And here we can see that this is indeed the case with the feature-rich molecular representations, ECFP6 and FCFP6 (each containing 2^32^ bits), being able to achieve an accuracy of above 98% with Naïve Bayes (comparable to levels attained by the other ML algorithms). Conversely, we should avoid using Naïve Bayes if the feature space for the molecular representation is small or sparse, like Estate (79 bits).

### Assessment of molecular representations for ligand-based screening

For all ML algorithms that we tested, the worst performing predictive models were derived from those that had used only the Estate fingerprint to represent the molecules. This is unsurprising considering that the Estate fingerprint is a 79-bit vector which serves as an atom-centred index that describes the arrangement/connectivity and electronic environment of atoms in a molecule^29,30^. However, what is surprising is that a relatively simple 79-bit molecular representation can achieve a predictive performance of above 85% accuracy on all but one ML algorithm – this result is almost on par with some Tox21 challenge winners, for example, dmlab which achieved an AUC-ROC of 0.828 for their screening of androgen receptor agonist and microsomes which achieved an AUC-ROC of 0.827 for their prediction of small molecule agonists of the estrogen receptor alpha (ER-alpha) signaling pathway^31,32^.

For predictive models using only a standalone fingerprint, ECFP6 consistently ranked among the top 3 fingerprints for all ML algorithms (Table 1). All models generated using ECFP6 as a standalone fingerprint managed to achieve an accuracy of above 96% with ECFP6 performing best with RF (98.48%) and SVM (98.93%). Furthermore, in all but one ML algorithm, the best performing models for these machine learners employed ECFP6 either in a paired fingerprint combination or as a standalone fingerprint. Even in the case of GBDT where the best model used a paired combination of AtomPair and Daylight-like fingerprints (97.36% accuracy), it should be noted that a model using a combination of ECFP6 with the AtomPair fingerprint followed closely with a 97.16% accuracy. Overall, the best predictive model was obtained from a paired combination of ECFP6 and Extended fingerprints in a SVM model (99.05%). Considering that ECFP6 occupies a feature space of 2^32^ bits, this fingerprint obviously provides a very detailed representation of a molecule and so, would undoubtedly be able to describe a molecule accurately and therefore, be useful in a classification task. These results suggest ECFP6 can be an effective molecular representation for LBVS regardless of whether it is used on its own or in combination with another fingerprint and would perform well irrespective of the ML algorithm in a data-driven approach.

So, is there any advantage in using a paired fingerprint combination to train our machines? As we can see with SVM, the pairing of Extended fingerprint with ECFP6 gave a slight improvement in accuracy and in fact, produced the best predictive model (Figure 1a and Supplemental Table 1). But more is not always better; there are instances where combining two molecular fingerprints offers no synergistic benefit and in fact, one might even observe a poorer performance from a paired fingerprint compared to the performance obtained by one of the standalone fingerprints that make up the pair. For example, the topological torsion fingerprint can achieve an average of 98.34% accuracy on its own but when combined with PubChem fingerprint, this drops to 97.86%. However, in some cases, this drop in accuracy may, in fact, offer an advantage; using ECFP6 on its own in the RF models, we obtain a 98.48% accuracy but pairing it with the Daylight-like fingerprint, this drops very slightly to 98.44%, but by sacrificing some accuracy, we attain near perfect precision for the model (increasing from 0.9892 to 0.9976) (Supplemental Table 1). For wet lab validation, compromising on accuracy to attain a high level of precision is desirable because it means that the chances of wasting time and resource validating a compound that is likely to be false positive is low.

**Figure 1a.**
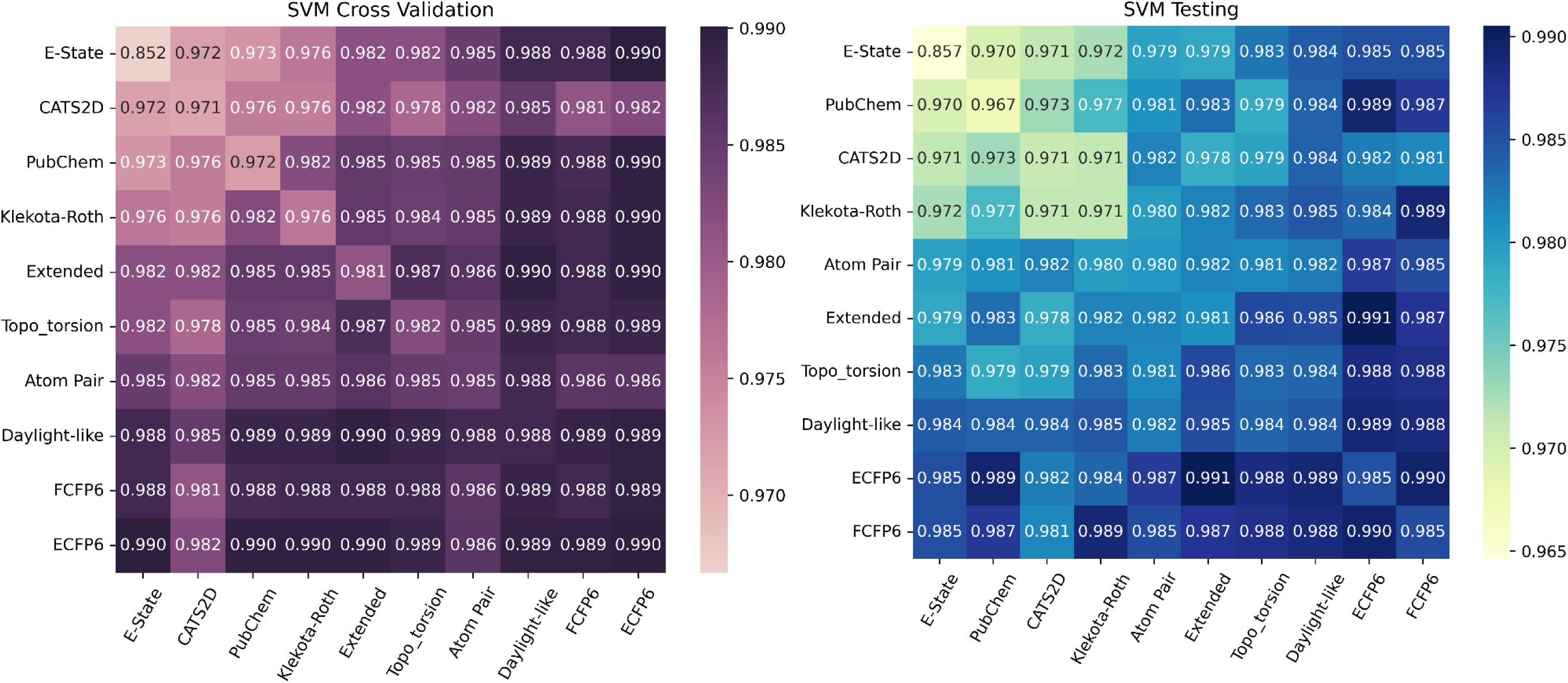
Accuracy of models generated with various single and paired molecular representations using support vector machine (SVM) during cross-validation (purple heatmap) and testing (blue heatmap)

**Table 1a:** The top performing standalone fingerprints for each of the 5 ML algorithms.

| Top 3 standalone fingerprints for each ML algorithm (% accuracy) |  |  |  |  |  |
| --- | --- | --- | --- | --- | --- |
| Rank | GBDT | NBayes | kNN | RF | SVM |
| 1. | AtomPair<br>(97.06) | ECFP6<br>(98.17) | FCFP6<br>(97.64) | ECFP6<br>(98.48) | FCFP6<br>(98.54) |
| 2. | Daylight-like<br>(96.6) | FCFP6<br>(98.06) | Extended<br>(97.36) | FCFP6<br>(98.24) | ECFP6<br>(98.5) |
| 3. | ECFP6 (96.46) | PubChem<br>(96.06) | ECFP6<br>(97.24) | Extended<br>(98.2) | Daylight-like<br>(98.44) |

**Table 1b:** The best and worst performing models using a merged fingerprint for all 5 ML algorithms.

|  |  | GBDT | NBayes | kNN | RF | SVM |
| --- | --- | --- | --- | --- | --- | --- |
| Molecular representations (% Accuracy) | Worst Performing Models | Estate<br>(88.19) | Estate<br>(59.44) | Estate<br>(85.13) | Estate<br>(92.13) | Estate<br>(85.67) |
|  | Best Performing Models | AtomPair + Daylight<br>(97.36) | [i] PubChem + FCFP6;<br>[ii] Extended + FCFP6<br>[iii] ECFP6 + FCFP6<br>(97.90) | FCFP6 + ECFP6<br>(98.22) | ECFP6<br>(98.48) | Extended + ECFP6<br>(99.05) |

### The role that dataset composition and size plays in performance

Current leading approaches in AI are data-hungry, requiring access to big data to accomplish its training tasks^33^. This may not work well in cheminformatics. For LBVS, we rely solely on prior knowledge of known ligands to train our models to predict potential hits, however, some targets have only a limited number of ligands. To illustrate this point, ChEMBL contained information for 6778 human protein targets at the time of writing (April, 2023) but more than half have 500 or less associated ligands^34^. Such targets may not benefit from model-centric approaches as the requisite large datasets are unavailable.

Although known ligands for a protein target is limited, the wealth of information in chemical databases like ChEMBL offer an almost unlimited opportunity to mine inactives for use in AI training for quantitative structure-activity relationship (QSAR) and LBVS. We wanted to see how the relative numbers of actives to inactives would impact performance and particularly, if an overabundance of inactives in the training dataset could compensate for a lack of actives on our four chosen predictive models: (i) SVM+ECFP6, (ii) RF+ECFP6, (iii) SVM+Daylight-like, and (iv) RF+Extended. We hypothesized that the use of a large dataset of inactives could improve performance when presented with a finite number of actives.

Model performance was tested in two different scenarios where: (1) the number of inactives increased equally with the number of actives in the training dataset and (2) the ratio of the number of inactives to actives increases for the training dataset. For this latter scenario, the number of inactives were fixed at 3600 while the number of actives were successively decreased from 3600 to 500 actives in the training dataset.

For the SVM models, when the number of actives is fixed, increasing the number of inactives did not have a significant impact on accuracy. With training datasets containing only 500 actives, increasing the number of inactives from 500 to 3600 resulted in a drop in the accuracy drop from 96.42% (Table 2b) to 95.73% (Table 2a) for the SVM+Daylight-like model - a difference of 0.69% in accuracy.

**Table 2a.**
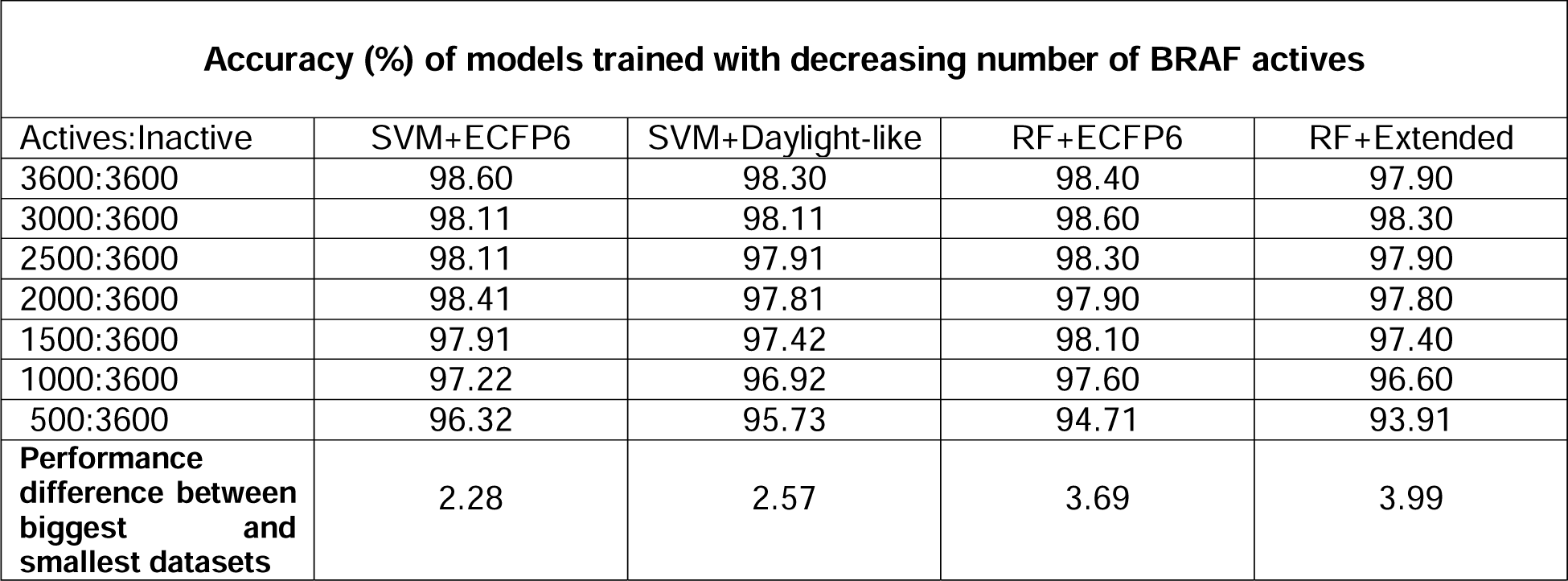
Accuracy (%) of models trained with an imbalanced training dataset where the number of BRAF actives is decreased but the number of BRAF inactives is maintained at a fixed number (3600)

**Table 2b.** Accuracy (%) of models trained with a balanced training dataset where the numbers of BRAF actives and BRAF inactives are both similarly decreased.

| **Accuracy (%) of models trained with decreasing number of BRAF actives and inactives** |  |  |  |  |
| --- | --- | --- | --- | --- |
| Actives:Inactive | SVM+ECFP6 | SVM+Daylight-like | RF+ECFP6 | RF+Extended |
| 3600:3600 | 98.60 | 98.30 | 98.40 | 97.90 |
| 3000:3000 | 98.31 | 98.01 | 98.40 | 98.10 |
| 2500:2500 | 98.21 | 97.91 | 98.20 | 97.30 |
| 2000:2000 | 98.11 | 97.81 | 98.00 | 97.50 |
| 1500:1500 | 98.01 | 97.62 | 98.20 | 97.30 |
| 1000:1000 | 97.62 | 97.12 | 97.60 | 96.00 |
| 500:500 | 97.22 | 96.42 | 97.20 | 96.10 |
| **Performance difference between biggest and smallest datasets** | 1.38 | 1.88 | 1.20 | 1.80 |

However, for the RF models, the difference in accuracy was conspicuous: the accuracy for the RF+ECFP6 model fell from 98.4% for the full training dataset (3600 actives, 3600 inactives) to 94.71% with the imbalanced (500 actives, 3600 inactives) training dataset. In fact, the performance of the model trained on a balanced (500 actives, 500 inactives) dataset still managed an accuracy of 97.2%. In other words, where the number of actives was fixed at 500, increasing the number of inactives from 500 to 3600 in the RF+ECFP6 model saw a drop in accuracy from 97.2% to 94.71% - a difference of 2.49%.

In short, our study shows that, given a finite number of actives, having a large number of inactives in the training dataset does not improve model accuracy. Furthermore, certain algorithms (like RF) are more sensitive to imbalanced training data than others (SVM) (Tables 2a and 2b) due to stronger affectations on the recall/sensitivity of RF models than SVM models (Table 2c).

**Table 2c.**
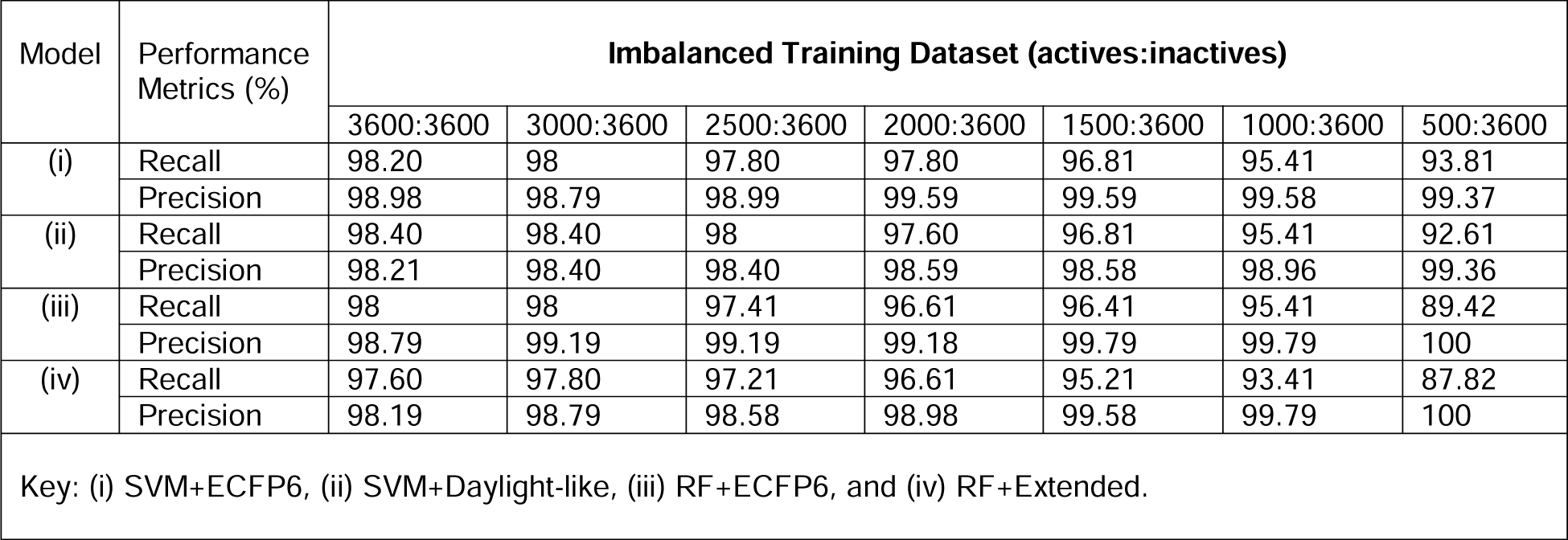
Recall and precision (%) of models trained with an imbalanced training dataset where the number of BRAF actives is decreased but the number of BRAF inactives is maintained at a fixed number (3600)

Regardless of the ML model and/or molecular representation used, recall/sensitivity decreases as the size of inactives in the training dataset increases (relative to the size of actives) while precision increases (Tables 2c and 2d). In a previous study where the number of actives were kept constant while increasing the number of inactives, Rodríguez-Pérez et al. had also found that precision improves as the ratio of inactives to actives increases ^35^. They also found that they needed at least 100 actives and 500 inactives to achieve near maximal recall (they achieved a mean recall of 87%) for their SVM model ^35^ but we found that the same cannot be said to be true for RF models – recall for RF models suffered greatly when the number of actives fall below 1000: for the RF+Extended model, recall fell from 93.41% when trained on (1000 actives, 3600 inactives) to 87.82% for the model trained on (500 actives, 3600 inactives) (Table 2c).

**Table 2d.**
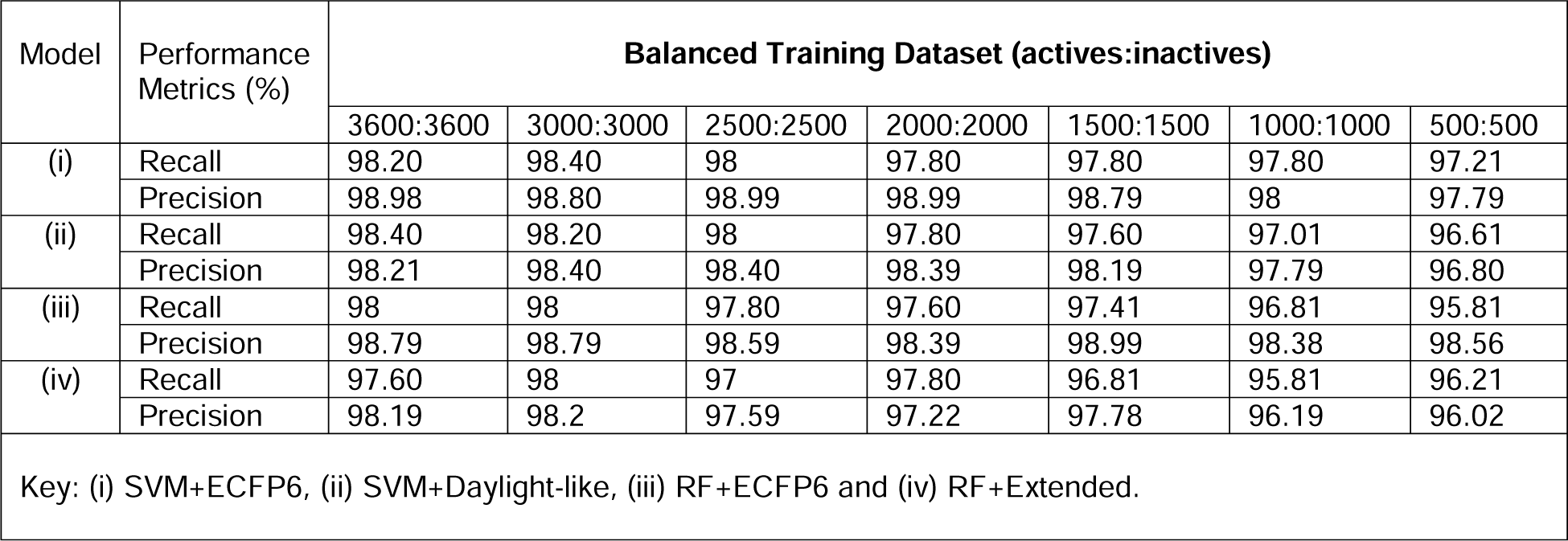
Recall and precision (%) of models trained with a balanced training dataset where the numbers of BRAF actives and BRAF inactives are both similarly decreased.

Although recall fell considerably, all SVM and RF models were able to achieve near-perfect (>99%) or perfect precision when we trained with only 500 actives but provided over seven times the number of inactives (3600 inactives) (Table 2c). It is also interesting to note that with a balanced training dataset, there was only a slight change in recall and precision for SVM and RF models even when both the number of actives and inactives in the training dataset drops from 3600 to 500, compared to models trained on the imbalanced datasets – in general, all models trained on a balanced dataset were able to attain >96% for both recall and precision (Tables 2c and 2d).

Despite the drop in recall and accuracy, we recommend that when faced with a limited number of actives for a target, one should increase the number of inactives used in training to lessen false positive predictions. It should be noted that no class weights were introduced when we trained our AI models on the imbalance datasets. Finally, the SVM+ECFP6 model was the most robust among the four models tested as its accuracy, recall and precision were little affected by the change in data composition or size.

### Choice of inactives is crucial for training

#### (i) Less active versus inactive

In many QSAR and LBVS studies, we often see the use of an arbitrary IC50 threshold to divide ligands into 2 classes, actives and inactives – for example, compounds that have an IC50 below 10 μM for a target protein are considered “actives” and those that have an IC50 of 10 μM or more are considered “inactives”^28,36–41^. We hypothesized that the use of such “inactive” ligands in training could reduce the model’s predictive performance since it is likely that these “inactives” are not really inactive but perhaps, just “less active”. We also recognise that the measurement of IC50 is dependent on the assay and condition under which it is tested. An IC50 curve shifts to the right if the expression/concentration of the protein is high in the cell/assay used to test the drug, thereby making what may be an “active” compound appear to be “inactive”. Here, we set out to see whether the use of these “less actives” affects a model’s predictive performance.

Before proceeding with our comparative studies with the “less actives”, we set out to first establish the baseline accuracy for our 4 predictive models. We created 10 new balanced, hold-out test sets to allow for an unbiased assessment of the 4 predictive models. The accuracy of each model is fairly consistent across all ten test sets, averaging at around 97% for the 4 predictive models (Table 3 and Supplemental Table 2). This suggests that the original test set used was not biased and that the assessment of the ML models in above study (Figures 1a-d) was valid. These results provide further evidence that the curated BRAF dataset used for training is of high quality and can be used as a reliable benchmark for future virtual screening methods. Finally, these results also form the baseline for the comparative studies below.

**Table 3.** Average accuracy for the ‘spiked-in’ “less active”-trained models based on testing with 10 balanced BRAF actives and inactives hold-out test sets.

| ML+Fingerprint | Average Accuracy(%) |  |  |  |
| --- | --- | --- | --- | --- |
|  | Training sets |  |  |  |
|  | 3600 actives /<br>3600 inactives | 3600 actives /<br>3420 inactives /<br>180 less actives | 3600 actives /<br>3240 inactives /<br>360 less actives | 3600 actives /<br>3060 inactives /<br>510 less actives |
| SVM+ECFP6 | 97.16 | 96.83 | 95.86 | 95.66 |
| SVM+Daylight-like | 97.17 | 96.40 | 95.83 | 94.91 |
| RF+ECFP6 | 97.74 | 96.24 | 95.81 | 95.34 |
| RF+Extended | 96.85 | 95.54 | 94.99 | 94.76 |

**Figure 1b.**
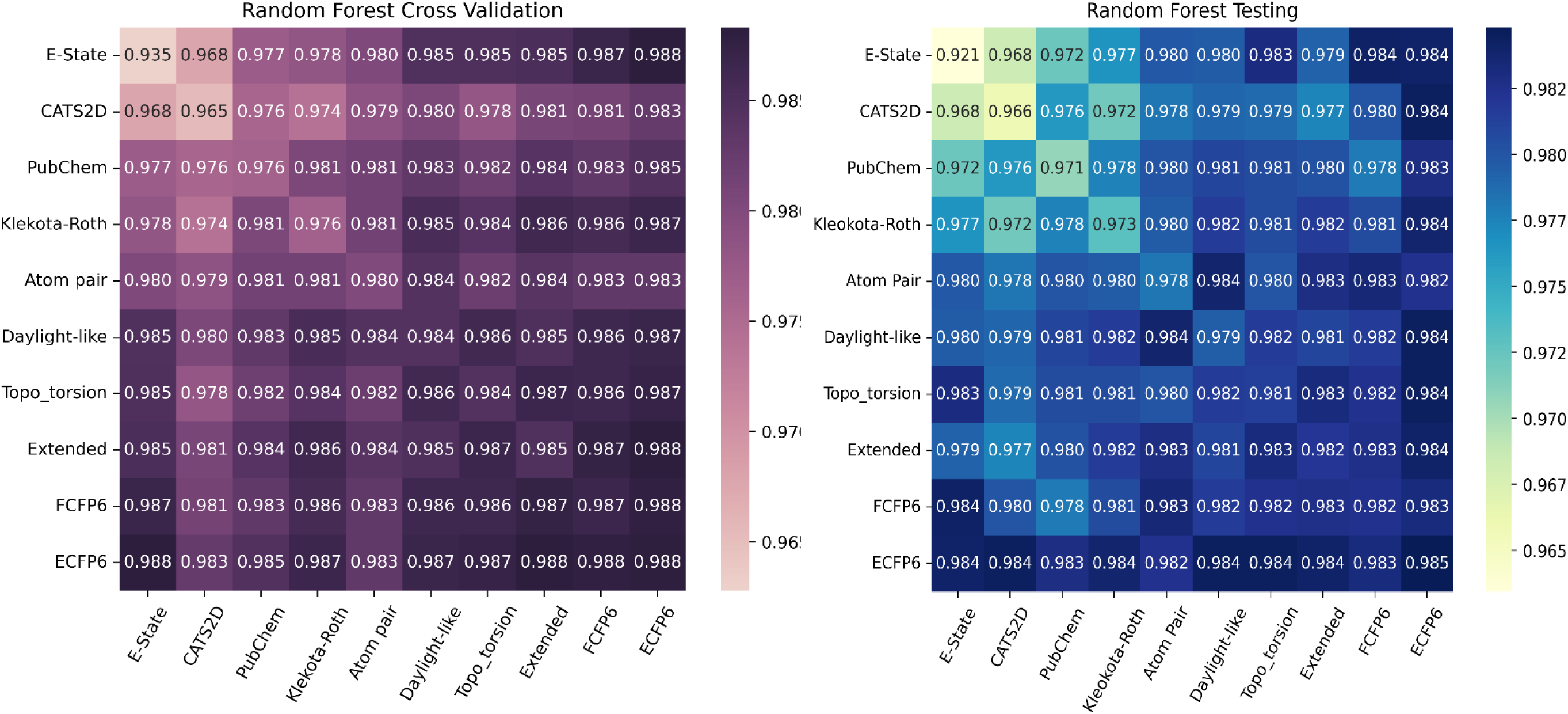
Accuracy of models generated with various single and paired molecular representations using random forest (RF) during cross-validation (purple heatmap) and testing (blue heatmap)

**Figure 1c.**
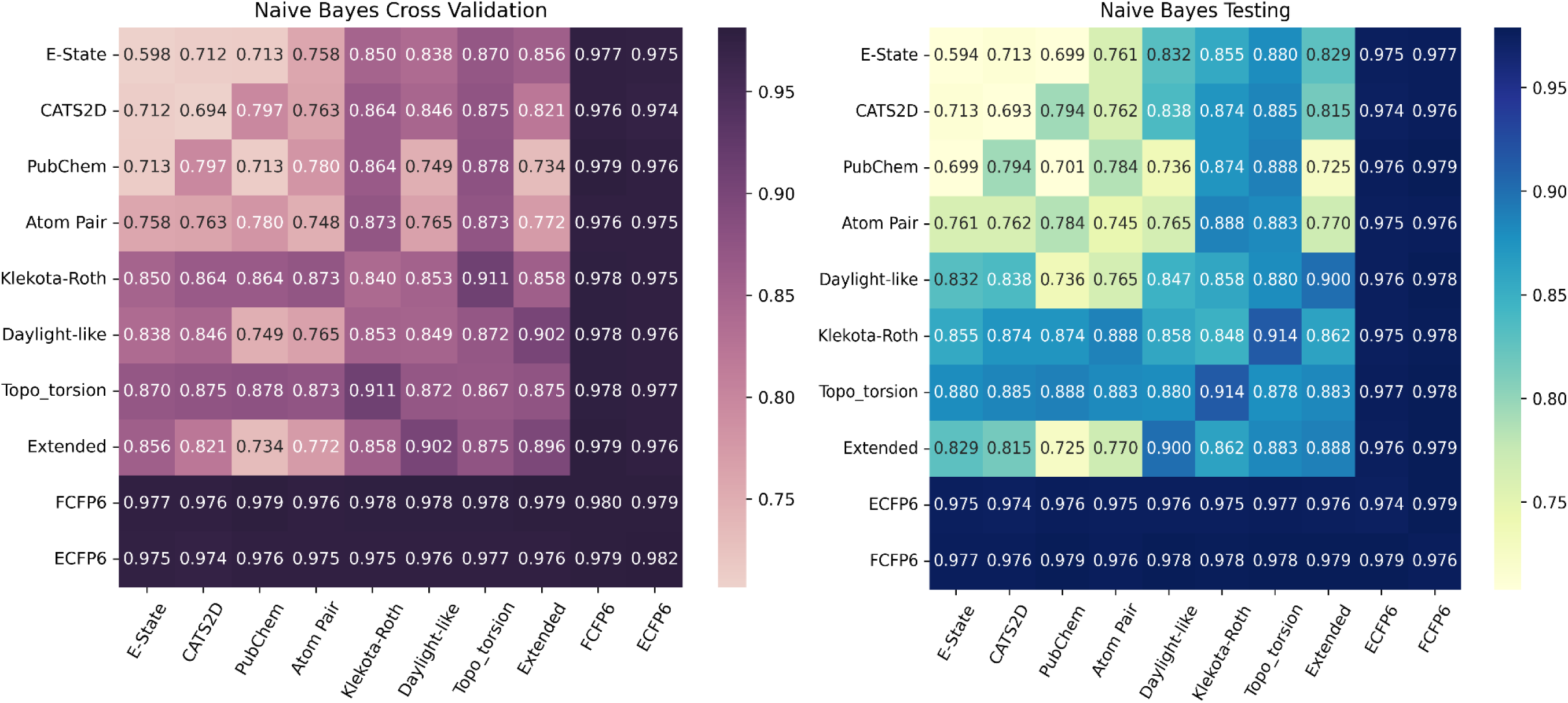
Accuracy of models generated with various single and paired molecular representations using naïve bayes (NBayes) during cross-validation (purple heatmap) and testing (blue heatmap)

**Figure 1d.**
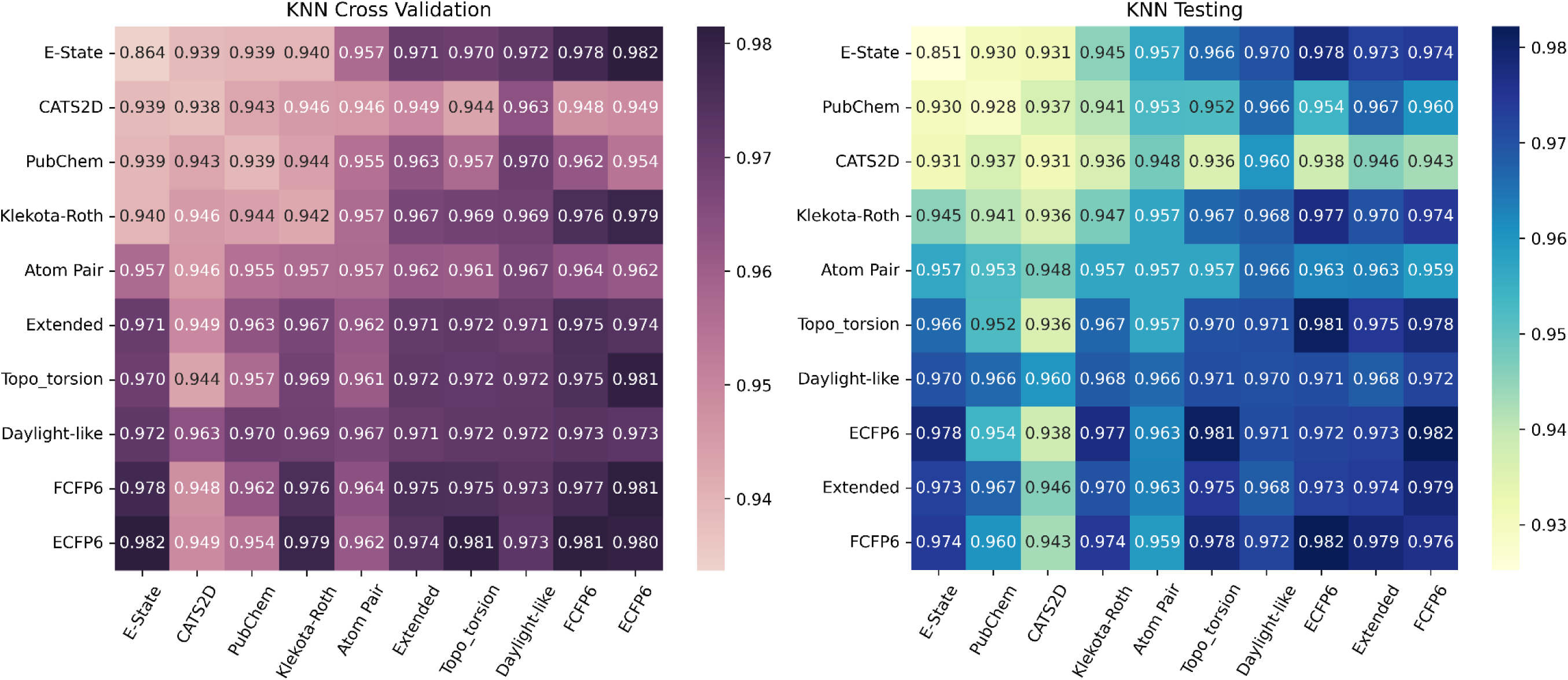
Accuracy of models generated with various single and paired molecular representations using k-nearest neighbour (kNN) during cross-validation (purple heatmap) and testing (blue heatmap)

**Figure 1e.**
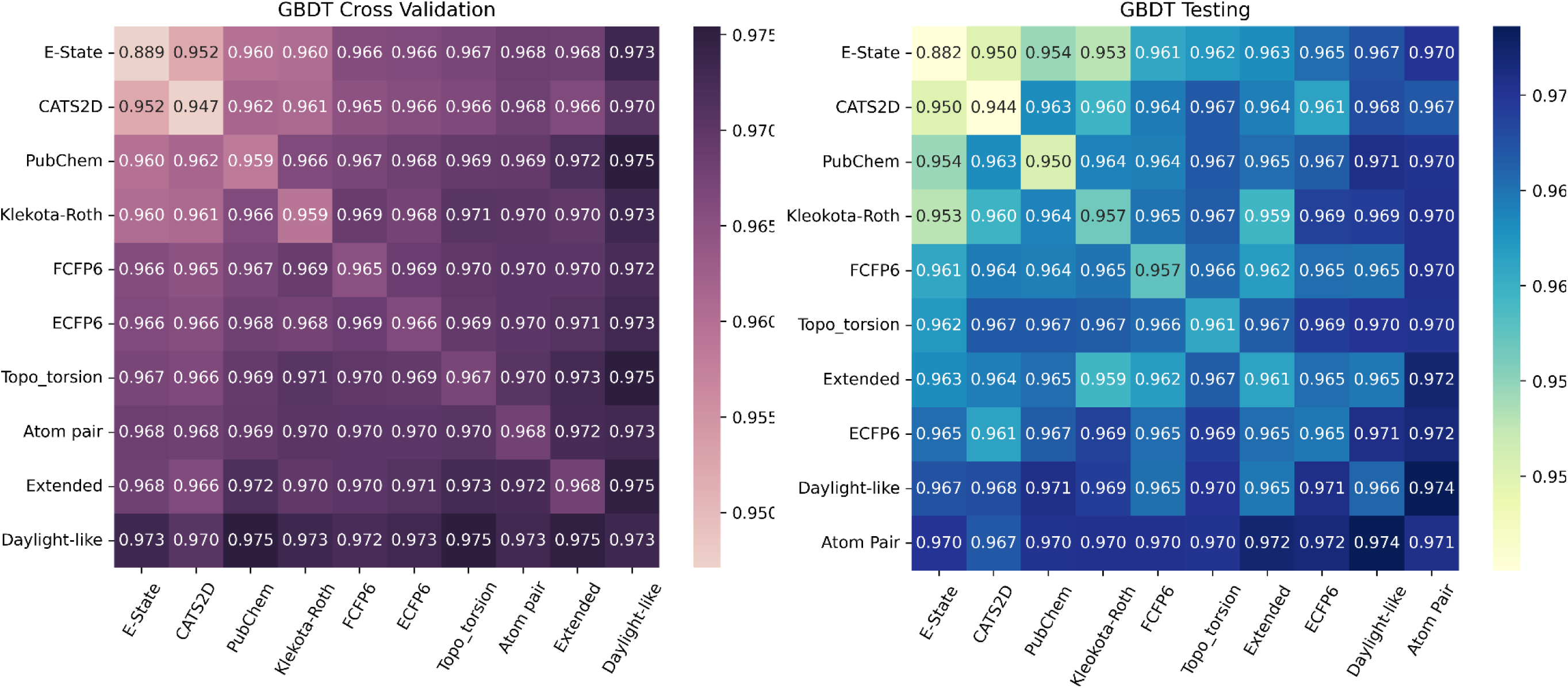
Accuracy of models generated with various single and paired molecular representations using gradient-boosting decision tree (GBDT) during cross-validation (purple heatmap) and testing (blue heatmap)

To test how “less actives” compounds would affect the predictive performance of the models, we identified 510 BRAF ligands in ChEMBL whose IC50 were more than 10 μM. We used these in a “spike-in” experiment to see if the introduction of these “less actives” in the original training dataset for our ML models would change their performance. We created 3 training datasets in which 5% (180), 10% (360) and 14% (510) of the inactives in the original training dataset were replaced with these “less actives”. In every instance, our results show that the level of accuracy of the model falls as the number of “less actives” in the training dataset increases (Table 3 and Supplemental Table 2). Not surprisingly, this drop in performance is due in large part to a drop in recall, meaning that the models trained on “less actives” are less able to correctly recognise certain actives (true positives) in the test set, probably because some of these actives share certain similarity to the “less actives” which were labelled as “inactive” in the training dataset. This demonstrates that the design and composition of the input data is important.

#### (ii) Using DUD-E decoys as inactives

The DUD-E benchmark dataset is commonly used for the development of SBVS methods^42^. Briefly, in DUD-E, a diverse set of actives is built for a target protein. Each active molecule is paired with a set of property matched decoys (PMD). PMDs are selected to be similar to each other and to known actives with respect to certain physico-chemical descriptors (e.g., molecular weight, hydrogen bond donors and acceptors) while being topologically dissimilar. It is believed that topological dissimilarity supports the assumption that the decoys are likely to be inactives because they are chemically different from the actives. The DUD-E database has 9,950 decoys for BRAF. Although originally developed for molecular docking screens, DUD-E’s decoys are also used as inactives in LBVS^43–48^. A recent study showed that the superior enrichment efficiency in convolutional neural network models for SBVS was achieved in the DUD-E dataset because of hidden bias in this dataset rather than a successful generalization of the pattern of protein-ligand interactions^49^. Therefore, could this hidden bias also have an impact on LBVS?

As with our spike-in studies with “less actives”, we wanted to see how the introduction of some decoys in training might affect model performance. Therefore, we selected at random 510 BRAF decoys and carried out the same spike-in experiments as described for the “less actives” experiment above. Unlike our models trained on datasets with “less actives” spiked-in, our models trained on datasets with decoys spiked-in do not show much difference in accuracy from our baseline models trained without any decoys. For example, for the RF+ECFP6 model trained on a dataset where 14% of the inactives were replaced with 510 decoys, the average accuracy on the 10 test datasets dropped by a mere 0.09% when compared with the baseline model (Table 4A); this is a stark contrast to the nearly 2% drop in accuracy when we used a dataset spiked-in with 510 “less actives” (Table 3). This is probably because decoys, unlike the “less actives”, are in fact topologically dissimilar from the BRAF actives and so, the use of some decoys to represent inactives does not severely compromise the performance of the model. However, the results for models trained on <u>only</u> actives and decoys was surprising.

**Table 4A.** Average accuracy for the ‘spiked-in’ decoy-trained models based on testing with 10 balanced BRAF actives and inactives hold-out test sets.

| ML+Fingerprint | Average Accuracy(%) |  |  |  |  |
| --- | --- | --- | --- | --- | --- |
|  | Training sets |  |  |  |  |
|  | 3600 actives /<br>3600 inactives | 3600 actives /<br>3420 inactives<br>/ 180 decoys | 3600 actives /<br>3240 inactives<br>/ 360 decoys | 3600 actives /<br>3060 inactives<br>/ 510 decoys | 3600 actives /<br>3600 decoys |
| SVM+ECFP6 | 97.16 | 96.95 | 96.85 | 96.80 | 93.85 |
| SVM+Daylight-like | 97.17 | 97.34 | 97.33 | 97.29 | 94.59 |
| RF+ECFP6 | 97.74 | 97.65 | 97.66 | 97.65 | 94.98 |
| RF+Extended | 96.85 | 96.49 | 96.69 | 96.79 | 94.09 |

For ML models trained on 3600 actives plus 3600 decoys, their accuracy when tested on a balanced test dataset of 1000 BRAF actives and decoys is near-perfect (above 99% for all ML models) (Table 4B; Supplemental Table 3). However, these models gave an average accuracy of around 94% when we tested it against our 10 balanced test datasets of 1000 BRAF actives and inactives (Table 4A; Supplemental Table 3). The drop in accuracy for models trained on 3600 BRAF actives plus 3600 decoys on our 10 test datasets is due to a fall in precision rather than recall. While recall for these models are above 98%, their precision drops by about 5% when compared with their corresponding baseline model. This means that models trained on only actives and decoys tend to give more false positive predictions. Conversely, when we tested our original ML models trained on our training dataset of 3600 BRAF actives plus 3600 BRAF inactives, we found that they gave an accuracy ranging from 96.85 to 97.74% on our 10 test datasets of BRAF actives and inactives but performed even better on the BRAF actives plus decoys test set, achieving near-perfect accuracy of 99% (Tables 4A & 4B; Supplemental Table 3). This is expected when we consider the fact that decoys are topologically dissimilar to BRAF actives, therefore, making it easy for our original ML models to discern between the BRAF actives and decoys. In summary, training with decoys is not recommended because it would result in a model that has a higher false positive rate. Testing with a dataset containing decoys is also not advisable because it may give an over-optimistic assessment of the predictive performance of the model.

**Table 4B.** Accuracy for the ‘spiked-in’ decoy-trained models based on testing with a balanced BRAF actives and decoys hold-out test set.

| ML+Fingerprint | Accuracy(%) |  |  |  |  |
| --- | --- | --- | --- | --- | --- |
|  | Training sets |  |  |  |  |
|  | 3600 actives /<br>3600 inactives | 3600 actives /<br>3420 inactives /<br>180 decoys | 3600 actives /<br>3240 inactives /<br>360 decoys | 3600 actives /<br>3060 inactives /<br>510 decoys | 3600 actives /<br>3600 decoys |
| SVM+ECFP6 | 99.00 | 98.80 | 98.80 | 98.80 | 99.30 |
| SVM+Daylight-like | 99.20 | 98.80 | 98.80 | 98.80 | 99.10 |
| RF+ECFP6 | 99.00 | 99.00 | 99.10 | 98.90 | 99.20 |
| RF+Extended | 98.90 | 98.50 | 98.80 | 98.70 | 99.10 |

## Discussion

“*Garbage in, garbage* out” or GIGO, is a concept familiar to all computer scientists denoting the importance and value of data quality in predictive modelling. But despite this wide acknowledgement, prevailing AI approaches tend to be model-centric, focusing more on the sophistication of the AI/ML architecture than the input. Such approaches may not pay enough heed towards resolving issues of data quality, data representativeness (not to be confused for data representations which is investigated in this study), and perhaps even feature-compatibility with the AI/ML models used. By adopting a model-centric AI approach, one hopes to tweak and improve the AI algorithms to improve its predictive performance while also enhancing model explainability or justifiability.

To pursue a data-centric AI approach, Dr Ng advocated developing a systematic approach to mine and clean large volumes of data in order to produce accurately-labelled data which is of high quality, consistent and error-free^7,8^. Data-centric AI appear to be gaining traction in the AI community^9,10,50,51^. It is also important that we do not confuse data-centric AI for data-driven AI: data-centric AI is an emerging science that studies techniques to improve the quality of datasets (that will be used to train the AI/ML models), whereas, data-driven AI is simply a paradigm that encourages the analysis of large datasets to improve AI performance. Many AI researchers have written to give a general perspective on the principles and critical considerations for the data construction process in data-centric AI ^9,10^ but to the best of our knowledge, there has been little or no attempts to determine and evaluate the parameters that are necessary for data construction in a data-centric AI approach for CADD. In fact, we can see that cheminformaticians still favour a model-centric AI approach when we look at studies published previously comparing the use of different ML or deep learning methods on different datasets ^27,28,37^. This emphasis on a model-centric AI approach for cheminformatics is also shared by authors of recent reviews surveying the current research landscape of CADD ^52–54^.

Whang et al. proposed that data acquisition, data labelling, data augmentation/improvement, data validation, data cleaning and data sanitization as key considerations in the practice of data-centric AI ^10^. However, we need to recognise that there are other considerations which are unique to chemistry and cheminformatics. A chemical’s property and structure need to be encoded in a meaningful manner so that a predictive AI can learn the salient features of a class of drugs for QSAR and virtual screening and this has given rise to various types of data representations for a molecule (that is, molecular descriptors and fingerprints). Thus, data representation is an important consideration that is somewhat unique to data construction for QSAR and virtual screening.

We believed that good data representation might be equally as important as high-quality data and so, we wanted to see how best to represent a molecule to the machine. In recent years, many newly created molecular representations have been proposed ^55–58^. However, instead of reinventing the wheel and creating another novel molecular representation, we tested to see if currently available molecular fingerprints are up to the task for virtual drug screening. Our study shows that certain molecular fingerprints are clearly superior and, in some cases, two molecular fingerprints can be combined to produce a performance which surpasses the performance of the individual molecular fingerprint, as seen when Extended and ECFP6 fingerprints were used with SVM. However, broadly speaking, there is very little difference in performance across different fingerprints. For example, looking at the RF models, most models trained using a single fingerprint could achieve a level of accuracy close to 97% or higher. The most surprising result came from the RF model using EState fingerprint which could achieve an accuracy of ∼92% - it is surprising because EState is only 79 bits but it was outperforming previously developed virtual drug screeners that used a more feature-rich molecular representation with deep learning methods ^28,59^. This suggests that current molecular fingerprints are more than up to the task for CADD if one can provide a high-quality dataset for training and testing.

Our study also demonstrates that one cannot pursue a data-centric AI approach to the exclusion of a model-centric approach because there are clearly certain AI models that perform better on certain AI task. Comparing between ML models, we see that although Estate fingerprint paired with RF achieves ∼92% accuracy, SVM with the same fingerprint achieves only 85% accuracy. This, however, does not suggest that SVM is not a good ML model for virtual screening because when given the “right” molecular representation (Extended and ECFP6 fingerprints), SVM can offer a predictive performance that surpasses all other predictive models that we tested. Thus, having the right model is just as important as having the right data and we cannot ignore the importance of a model-centric approach --- both model and data-centric approaches should be seen as complimentary than mutually exclusive. In future, in-depth studies to understand complementariness between models and data should be performed.

Interest in deep learning methods for QSAR and virtual screening have been on the rise following the success of Dahl et al.’s MT-DNN in the Merck Molecular Activity Challenge^2,3^ and subsequently, Mayr et al.’s DeepTox deep neural networks in the Tox21 Challenge^59^. Studies have shown that deep learning methods consistently outperform conventional ML methods for rational drug discovery^1^. For example, DeepTox (the overall winner of the Tox21 Challenge) which used a deep neural network for toxicity prediction could achieve AUC-ROCs of between 0.793-0.942 for 12 nuclear receptor (NR) and stress response (SR) pathway assays^59^. The best ligand-based screening model for DeepTox was achieved for ligands of the aryl hydrocarbon receptor (AhR) which gave an AUC-ROC of 0.928. However, we need to recognise certain limitations of deep learning methods. In comparison to conventional ML algorithms, deep neural networks require a much larger amount of data to ensure model generalizability and prevent overfitting. This presents a problem for LBVS because many druggable protein targets have very few known ligands. To illustrate this point, ChEMBL contains information for 6778 human protein targets (as of April, 2023) but more than half (∼3800) have 100 or less compounds/ligands associated with each protein^34^. For this reason, we chose to focus only on conventional ML algorithms and we show here that conventional ML models can give superior performance for virtual screening, without the need to resort to more data-hungry deep learning methods. This limitation of deep neural networks also makes it clear that alternative paradigms such as data-centricity, favouring quality over quantity, is more likely to be successful in novel drug discovery using AI/ML.

Until recently, it has been difficult to give a reliable assessment of what constitutes best practice for LBVS since there are no benchmark datasets that can produce ML models capable of giving near-perfect predictive performance. In the absence of a high-quality benchmark dataset, evaluating experiments such as the ones carried out here would be difficult because any observed change in performance could be attributed to the behaviour of an imperfect model rather than a reflection of the use of inappropriate data. We have now generated a high-quality benchmark dataset and were able to show that current practices and use of data lead to serious shortcomings in the state-of-the-art predictive models which, in turn, calls for a reassessment of these model’s true performance. Our work indicates that there is little or no shortcomings with either current predictive AI/ML algorithms or most molecular representations when used in LBVS because the differences in their performance are minimal. Ultimately, the biggest impact on predictive performance comes from data size, composition and poor data quality.

Through this work, we hope to use a single case study to demonstrate the power of data-centric paradigm. As this approach becomes more widespread and better understood across use cases, a foundation for data-centric AI approaches in cheminformatics could eventually be laid. With the exponential growth in chemical databases like ChEMBL and PubChem, it becomes increasingly difficult to manually curate a high-quality dataset when we are dealing with large numbers of compounds/ligands. It is therefore important for us to consider how data-centric AI approaches can help. We propose that pursuing data-centricity require first understanding the nature of our data across 4 pillars: data representation, data quality, data quantity, and data composition on a model’s predictive performance (i.e., what attributes or properties of these 4 pillars contribute to high-quality data for training and testing). If, for example, we do not know what type of data diminishes data quality, then how can we be expected to build a data-centric AI approach for cheminformatics? Our study demonstrates that manipulation of these pillars of data-centricity has affectation on model performance. Pending future generalization studies on more use cases, our work paves the way towards the establishment of a framework for a data-centric AI approach for virtual screening and CADD.

## Method

### Machine Learning Models

We tested the performance of five machine learning methods representing different strategies. These include, k-nearest neighbours (kNN), Naïve Bayes (NBayes), gradient-boosted decision tree (GBDT), random forest (RF) and support vector machine (SVM) (python ver. 9.1.2 / sklearn ver.1.0.2), using the BRAF dataset described below.

For each of the 55 molecular representations that we tested, we used 5 training datasets on the above machine learning methods. We did not perform any feature scaling or selection. Each training set underwent 10-fold cross-validation to optimize the hyperparameters for the SVM, kNN and NBayes models and to estimate the accuracy of the models. Grid search was used to determine the optimal values of the hyperparameters. For the RF and GBDT models, the default parameters were used with no additional tuning.

Accuracy, precision, and recall were used as performance metrics.

## Dataset

Instead of using a readily available benchmark dataset (such as the BACE1 inhibitors dataset by Subramanian et al.^28^), we manually curated and built a dataset of BRAF actives and inactives to ensure that we would have a high quality dataset for our study. All data was mined from ChEMBL in June/July 2022 (ChEMBL Release 30) ^34^. Actives were defined as validated BRAF ligands with an IC50 of less than 10 µM, while inactives were carefully selected based on the fact that they have no known pharmacological activity against BRAF. In total, we identified 4100 BRAF actives and around 24,000 compounds that we deemed to be inactive against BRAF. From this initial set of compounds, we randomly selected 3600 BRAF actives to be part of the training dataset with the remaining 500 actives becoming a part of the hold-out test set. To avoid training bias from any single curated dataset, we created 5 balanced training datasets, each containing the 3600 BRAF actives with an equal number of inactives and where each training dataset contained a unique set of inactives that was not shared, partially or wholly, with the other 4 training datasets. Unless there were some hidden biases in one or more of the training datasets, we expected the performances of the 5 trained models for each molecular representation to be similar. We also created a single balanced test set with 1000 BRAF actives and inactives that would be used to give a fair and objective comparison of the predictive models. (Please refer to Supplementary data (Part 1) for these training and test datasets)

We chose four top predictive models employing a single molecular fingerprint that had been trained with the third training dataset for the experiments investigating data size, data composition and data quality. The models chosen were: [i] SVM+ECFP6, [ii] SVM+Daylight-like, [iii] RF+ECFP6, and [iv] RF+Extended. The performance of these four original ML models formed the baseline for our comparative analyses and represented the optimal performance achievable for the respective model. Additionally, we generated new predictive models where [i] SVM+ECFP6, [ii] SVM+Daylight-like, [iii] RF+ECFP6, and [iv] RF+Extended were trained on smaller datasets to compare their performance against the baseline performance. The original training dataset was a balanced dataset of 7200 BRAF actives and inactives but the smaller training datasets composed of the following numbers of BRAF actives to inactives: (i) 3000:3600; (ii) 2500:3600; (iii) 2000:3600, (iv) 1500: 3600; (v) 1000:3600; (vi) 500:3600; (vii) 3000:3000; (viii) 2500:2500; (xi) 2000:2000, (x) 1500:1500; (xi) 1000:1000; (xii) 500:500. The actives and inactives in these smaller datasets were randomly selected to avoid any unintentional bias.

For this study, we generated 10 new balanced hold-out test datasets of 1000 BRAF actives and inactives to give us an unbiased gauge of the accuracy that our models can achieve. We define a compound as “inactive” if there are no known pharmacological assays for the said compound on our target, BRAF.

We found 510 BRAF ligands that have an IC50 of more than 10 μM in the ChEMBL database which we labelled as “less active” ^34^. These “less active” ligands were used for spike-in experiments where we replaced either 180 (5%), 360 (10%) or 510 (14%) inactives in our original training dataset with a similar number of “less active” ligands.

The DUD-E database contains 9,950 decoys for BRAF^42^. We randomly selected 3600 decoys to create a training dataset of 7200 BRAF actives and decoys. In addition, we also created a balanced hold-out test set of 1000 BRAF actives and decoys. As with the “less actives” spike-in experiments, we also created 3 separate training datasets that had either 180 (5%), 360 (10%) or 510 (14%) decoys replacing a similar number of inactives in our original training dataset of 3600 actives and 3600 inactives.

Please refer to Supplementary data (Part 2) for training and test datasets used in these studies.

### Fingerprint Generation

RDKit and the Chemistry Development Kit (CDK) were used to compute the fingerprints for the molecules in this study^60,61^. The ten standalone molecular fingerprints, namely,

1. Estate fingerprints (CDK)^29,30^
2. PubChem fingerprints (CDK)^11,62^
3. Klekota-Roth fingerprints (CDK)^63^
4. Extended-Connectivity fingerprints 6 (ECFP6) (CDK)^64,65^
5. Functional-Class fingerprints 6 (FCFP6) (CDK)^64,65^
6. Extended fingerprints (CDK)^66^
7. Topological Torsion fingerprints (topo_torsion) (RDKit)^67^
8. Atom Pairs fingerprints (RDKit)^68^
9. Daylight-like RDKit fingerprints (RDKit)^60^
10. CATS2D fingerprints (RDKit)^69^

and 45 paired combinations of these were used in this study.

## Supporting information

Supplementary Section - Additional Experiments

Supplementary data (Part 1)

Supplementary data (Part 2)

Supplementary data (Part 3)

## Acknowledgement

The authors wish to thank Dr. Cameron Osborne for taking time to read our manuscript and offer valuable comments and suggestions to improve its readability.

This research/project is supported by the National Research Foundation, Singapore under its Industry Alignment Fund – Pre-positioning (IAF-PP) Funding Initiative. Any opinions, findings and conclusions or recommendations expressed in this material are those of the author(s) and do not reflect the views of National Research Foundation, Singapore.

WWBG acknowledges support from a Ministry of Education (MOE), Singapore Tier 1 and SUG grant (Grant No. RS08/21).

