## Supplementary Section - Additional Experiments for "Establishing the foundations for a data-centric AI approach for virtual drug screening through a systematic assessment of the properties of chemical data"

The aim of this supplementary section is to provide further proof of the robustness of our data-centric AI approach for virtual drug screening and address the concerns raised by the reviewers and editors.

**Impact of training dataset size on predictive performance**

ChEMBL contained information for 6778 human protein targets but slightly more than half have 500 or less associated ligands^1^. Thus, we wanted to show the level of performance that could be achieved with our approach if we trained with 500 actives and 500 inactives (Table 2b). We demonstrated that both support vector machine (SVM) and random forest (RF) models using ECFP6 could attain an accuracy of at least 97.2%.

Here, we want to show how further decreasing the training dataset size affects various performance metrics. Since both SVM and RF models performed equally well using ECFP6 (Table 2b), we chose to study the impact of a decreasing training dataset size on the RF model in this instance.

We created training datasets of decreasing size, consisting of:

1) 250 BRAF actives & 250 BRAF inactives
2) 100 BRAF actives & 100 BRAF inactives
3) 50 BRAF actives & 50 BRAF inactives
4) 10 BRAF actives & 10 BRAF inactives

The models trained on the above datasets were tested on the hold-out test dataset and the results are shown in Table SS1.

We can see that with only as little as 100 BRAF actives and 100 BRAF inactives, our RF model can still give an exceedingly high level of accuracy (above 90%) (Table SS1). Dropping below 100 BRAF actives had a profound effect on the model’s recall (which fell from over 90% to 78%). This was also previously observed by Rodríguez-Pérez et al. using SVM ^2^. Thus, we would recommend to have at least 100 actives/100 inactives in the training dataset in order to maintain a reasonable level of predictive performance.

**Further proof to support our initial work**

We created 10 new balanced hold-out test sets to demonstrate that performance originally obtained by on the first test set had not been unknowingly biased such as to allow our predictive models to achieve near-perfect accuracy. The results on these 10 balanced hold-out test sets are given in Supplemental Table 2 and summarized in Table 3. We can see that the average accuracy obtained with the RF model using the ECFP6 fingerprint was 97.74% on the 10 test sets. This compared well with the result of the original hold-out test set which gave an accuracy of 98.48% for the same RF model using the ECFP6 fingerprint (Table 1a). Despite this, there were concerns raised by the reviewers that maybe active and inactive molecules are very dissimilar and therefore, any ML algorithm and fingerprint could have been used to obtain good prediction results. However, one needs to recognize that ligand-based virtual screening works on the basis of the Chemical Similarity Principle that states that structurally similar compounds tend to possess similar properties and biological activity. Thus, it would be expected that the BRAF actives are very similar to each other while being very dissimilar to other chemical compounds. However, we wanted to show that the performance of our AI models is indeed the result of our superior data-centric approach and not because an inherent characteristic of the BRAF ligands that gave our AI model a biased advantage.

(i) **Comparing the degree of similarity/dissimilarity of BRAF actives and inactives**

Here, we wanted to show just how similar or dissimilar BRAF ligands were to each other and to other compounds/inactives. Thus, we calculated the Tanimoto coefficients between

a) BRAF active versus BRAF active: that is, calculating the Tanimoto coefficient of all 4101 BRAF actives against each other
b) BRAF active versus test set: that is, calculating the Tanimoto coefficient of 4101 BRAF actives against the 1000 compounds (500 BRAF actives and 500 BRAF inactives) in the original hold-out test set used to ascertain the performance of the predictive models given in Figures 1a-e
c) BRAF active versus inactive: that is, calculating the Tanimoto coefficient of 4101 BRAF actives against all BRAF inactives used in our training and testing
d) BRAF active versus ChEMBL: that is, calculating the Tanimoto coefficient of 4101 BRAF actives against all compounds in ChEMBL

Since the ECFP6 fingerprints were used for our RF models, we used ECFP6 when calculating the Tanimoto coefficients for the above comparisons. Figure SS1 gives the distribution plot of the Tanimoto coefficients for the four conditions.

The Tanimoto coefficient measures the degree of similarity between two compounds and its value ranges from 0 to +1 (where +1 denotes the highest similarity possible between two compounds). The results show that the BRAF actives are, in fact, very dissimilar from the BRAF inactives used in our study with a mean Tanimoto coefficient of only 0.0905 and it is also shown to be highly dissimilar when compared against 2 million-odd ChEMBL compounds, giving a mean Tanimoto coefficient value of 0.0682. Thus, it may appear that perhaps BRAF actives are indeed highly dissimilar to other types of compounds and so, that would make it easy for a predictive machine to differentiate BRAF ligands from other non-BRAF ligand compounds. However, a closer look when BRAF actives are compared among themselves, we see that the mean Tanimoto coefficient is also very low (mean = 0.1134; std. dev.= 0.086). This means that BRAF ligands don’t share a lot of similarity that might help group them into a class and help delineate them from other molecules. This shouldn’t come as a surprise since ECFP6 is a 2^32^ -bit fingerprint ^3,4^ and therefore, for any two compounds to share a high degree of similarity in ECFP6, say for example, a Tanimoto coefficient of 0.8 or more, they would have to have more than 3 billion features in common! Comparing the mean Tanimoto coefficient calculated for (i) BRAF actives versus compounds in the original hold-out test set (mean=0.07; std. dev.= 0.026) with that of (ii) BRAF actives versus all ChEMBL compounds (mean=0.0682; std. dev.= 0.025), we find that the degree of dissimilarity between BRAF actives and inactives in our original test set is equivalent to that found between BRAF actives and inactives in ChEMBL. This therefore shows that the performance of our AI models did not perform well because of a high degree of similarity of within the class of BRAF ligands and/or high degree of dissimilarity between BRAF actives and inactives.

(ii) **Performance in a real world screening**

We decided to use a RF model trained on 1000 BRAF actives and 1000 BRAF inactives to demonstrate how our model might perform in a real world screening of a large, diverse chemical dataset. We showed that this model was able to achieve an accuracy of 97.6% in a balanced test set (see Table 2b). However, here, our hold-out test set consisted of 3101 BRAF actives plus all ChEMBL compounds as inactives. The results of this screening are shown in Table SS3.

Testing our model with a balanced hold-out test set gave us an accuracy of 97.6% (Table 2b) but when our model is tested against a large diverse, imbalanced test set, we found that it could still attain an accuracy of 97% and a recall of 97.07%. Understandably, precision (4.06%) dropped appreciably due the large numbers of false positives but this is to be expected when one is screening a large dataset.

(iii) **Our approach can be replicated**

To further prove that our novel approach does not inordinately favour the search for BRAF ligands, we developed a virtual screening platform for β‑Secretase 1 (BACE-1) inhibitors. A benchmark dataset of β‑Secretase 1 (BACE-1) inhibitors was previously developed by Subramanian *et al*. ^5^. This dataset was developed for ligand-based approaches and has been widely cited (168 times). The authors had previously reported that they could achieve a classification accuracy of 76% for a Bayesian model using the ECFP6 fingerprint, with recall reaching 87% while their RF model using the Canvas constitutional, physicochemical, and topological descriptors was reported to achieve 81% accuracy and a recall of 77%. The precision of their models was not given in their paper but based on the available data in the paper, we were able to calculate the precision of their top three predictive models in order to give a comparison of our model with theirs (Table SS4).

Following the same criteria that we used in creating our BRAF ligands dataset, we selected 1273 BACE-1 inhibitors with an IC50 of less than 10 µM from the Subramanian *et al*. dataset to be our actives. We also selected an equal number of compounds from ChEMBL with no known pharmacological activity against BACE-1 as inactives. In total, our new dataset has 1595 actives and 1595 inactives. We did an 80:20 train-test split on this new dataset and results for an RF model using ECFP6 are presented in Table SS4. As with our RF predictive model for BRAF ligands, the model built for BACE-1 inhibitors was also able to reach near-perfect accuracy (98.59%) on the hold-out test set and achieved exceedingly high recall (97.81%) and precision (99.36%). Our RF + ECFP6 model trained on our newly created dataset outperformed all predictive models built by Subramanian *et al*. ^5^ There is also no overfitting of our model since there is minimal difference in performance between the cross-validation and test. Furthermore, we demonstrate that in a large diverse test dataset where known BACE-1 inhibitors are combined with other ChEMBL compounds, the accuracy is exceptionally high (99.61%), while retaining a recall of 97.18% (Table SS4b). Please refer to Supplementary data (Part 3) for training and test datasets used in these studies.

We believe that we have demonstrated conclusively that our approach can be applied to other classes of drugs and had not performed well with BRAF ligands because of a bias that may have arisen due to an unexplainable feature of BRAF ligands that might make them easily distinguishable from other compounds. Furthermore, our approach is robust and can still attain a high degree of accuracy even when the training dataset is small.

Table SS1: Accuracy, recall and precision (%) of Random Forest models trained with a balanced training dataset with decreasing number of BRAF actives and inactives

| Actives:Inactives | True Negative (TN) | False Positive (FP) | False Negative (FN) | True Positive (TP) | Accuracy (%) | Recall (%) | Precision (%) |
| --- | --- | --- | --- | --- | --- | --- | --- |
| 250:250 | 484 | 16 | 21 | 480 | 96.30 | 95.81 | 96.77 |
| 100:100 | 483 | 17 | 45 | 456 | 93.81 | 91.02 | 96.41 |
| 50:50 | 490 | 10 | 106 | 395 | 88.41 | 78.84 | 97.53 |
| 10:10 | 459 | 41 | 110 | 391 | 84.92 | 78.04 | 90.51 |

Table SS2: Statistical calculations of mean and standard deviations calculated for the 4 distribution plots in Figure SS1

| Condition | Tanimoto Coefficient | |
| --- | --- | --- |
|  | Mean | Standard Deviation |
| (a) BRAF active v BRAF active | 0.1134 | 0.086 |
| (b) BRAF active v Test set | 0.07 | 0.026 |
| (c) BRAF active v Inactive | 0.0905 | 0.066 |
| (d) BRAF active v ChEMBL compounds | 0.0682 | 0.025 |

Table SS3: Performance of our RF model for BRAF ligands in a screening of a large and diverse chemical dataset

| Test set | True Negative (TN) | False Positive (FP) | False Negative (FN) | True Positive (TP) | Accuracy  (%) | Recall (%) | Precision (%) |
| --- | --- | --- | --- | --- | --- | --- | --- |
| 3101 BRAF actives plus 2,367,573 ChEMBL compounds | 2296519 | 71054 | 91 | 3010 | 97.00 | 97.07 | 4.06 |

Table SS4a: The table shows the performance of: ^a^ the Random Forest (RF) model using ECFP6 developed with our BACE-1 ligand dataset and ^b^ the top three predictive models developed by Subramanian *et al*. Performance metrics for accuracy and recall for Subramanian *et al*.’s three models were obtained from Table 1 of their paper, however, ^c^ the precision of their models was calculated by us based on the available data given in their paper.

| Model | Cross-validation | | | Test set | | |
| --- | --- | --- | --- | --- | --- | --- |
|  | Accuracy (%) | Recall  (%) | Precision (%) | Accuracy (%) | Recall  (%) | Precision (%) |
| RF +  ECFP6 ^a^ | 98.43 | 97.38 | 99.46 | 98.59 | 97.81 | 99.36 |
| Bayes + ECFP6 ^b^ | 79 | 90 | 70.55 ^c^ | 76 | 87 | 67.72 ^c^ |
| RF + Canvass ^b^ | 99 | 99 | 98.74 ^c^ | 81 | 77 | 79.29 ^c^ |
| DNN +  Canvass ^b^ | 88 | 85 | 87.47 ^c^ | 82 | 75 | 82.56 ^c^ |

Table SS4b: This table shows the performance of our Random Forest (RF) model using ECFP6 on a hold-out test set comprising of 319 BACE-1 ligands (actives) and all compounds not identified as BACE-1 ligands in ChEMBL (inactives)

| Model | True Negative (TN) | False Positive (FP) | False Negative (FN) | True Positive (TP) | Accuracy  (%) | Recall (%) | Precision (%) |
| --- | --- | --- | --- | --- | --- | --- | --- |
| RF +  ECFP6 ^a^ | 2363816 | 9177 | 9 | 310 | 99.61 | 97.18 | 3.27 |

Figure SS1: Distribution plots of the Tanimoto coefficients calculated using the ECFP6 fingerprint for four conditions: (a) BRAF active versus BRAF active, (b) BRAF active versus compounds in the hold-out test set, (c) BRAF active versus all BRAF inactives that were used (for both training and testing), and (d) BRAF active versus all ChEMBL compounds

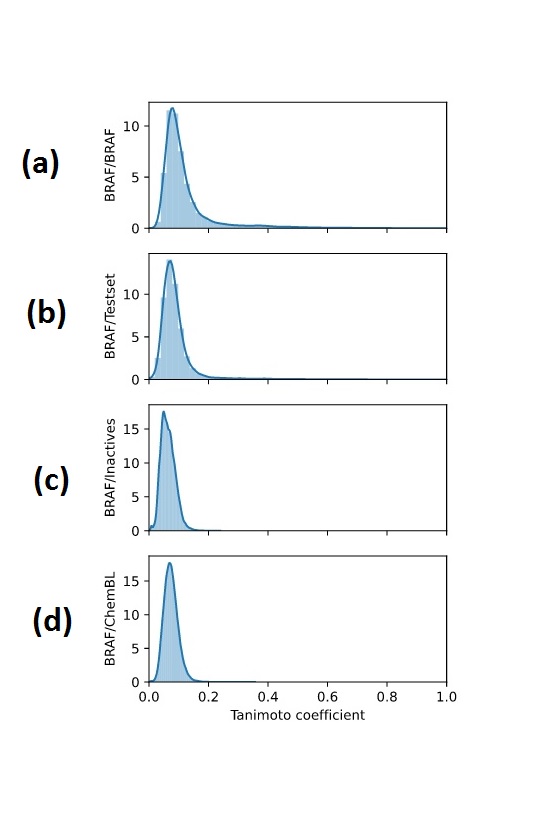
